# Multi-omics analysis reveals vitamin D metabolism, hyper-IgE genes, and epithelial barrier dysfunction in hazelnut allergy

**DOI:** 10.64898/2025.12.11.692994

**Authors:** Alexander C.S.N. Jeanrenaud, Aleix Arnau-Soler, Ahla Ghauri, Ingo Marenholz, Philipp Mertins, Kirsten Beyer, Margitta Worm, Young-Ae Lee

## Abstract

**Background:** Hazelnut allergy is a major cause of food-induced anaphylaxis yet remains poorly defined at the molecular level.

**Objective:** We aimed to identify molecular differences between individuals with primary hazelnut allergy and nonallergic controls by investigating a comprehensive spectrum of omics profiles in immune cells.

**Methods:** We analysed DNA methylation, transcriptomic and proteomic profiles in hazelnut-stimulated and unstimulated immune cells.

**Results:** Across analyses, we identified 80 differentially methylated signatures, 125 differentially expressed genes, and 11 differentially secreted proteins associated with hazelnut allergy. DNA methylation signatures were highly concordant between unstimulated and stimulated conditions, consistent with stable epigenetic remodelling. Key findings implicated *ZNF341*, associated with a rare monogenic hyper-IgE syndrome, and *ARL2*, both linked to STAT3-mediated IgE dysregulation. Additionally, we identified a differentially methylated region (DMR) overlapping the *T Helper Type 2 Locus Control Region Associated RNA* (*TH2LCRR*) in the cytokine gene cluster, suggesting an epigenetic mechanism contributing to IL-5 and IL-13 upregulation. Antigen stimulation was required to reveal hazelnut-specific transcriptional and proteomic signals. Integration of these data demonstrated that IL-5 expression could distinguish both groups. We identified signals in epithelial barrier genes of the gut and skin (*TRIM31*, *TRIM40*, *CDSN*), activation of the vitamin D pathway (*CYP27B1*, *IL32*), and nominate additional signals (*PHACTR1, MFHAS1, SPRED2, GALNT5/GALNTL4, NSMCE1-DT*) for follow-up.

**Conclusion:** Our study confirms type-2 cytokines, FcεRI, and JAK–STAT signalling and uncovers novel links to monogenic hyper-IgE syndrome, activation of vitamin-D pathways, and gut/skin barrier genes, yielding a catalogue of candidate biomarkers for mechanistic studies and prospective validation.

**Key messages:**

- Antigen-specific multi-omics analysis confirms JAK-STAT and Th2 control pathways, with IL-5 emerging as a key marker distinguishing hazelnut-allergic from nonallergic individuals.
- Epigenetic and transcriptomic analysis points to roles for vitamin D metabolism, hyper-IgE-associated genes, and epithelial barrier dysfunction in hazelnut allergy
- This first antigen-specific methylation and multi-omics discovery study in hazelnut allergy provides candidate pathways and genes to guide future studies

**Capsule summary:** This antigen-specific multi-omics study confirms JAK-STAT/Th2 control pathways, pinpoints the IL-5 response as biomarker distinguishing hazelnut allergy, implicates vitamin D metabolism, hyper-IgE genes, and epithelial barrier dysfunction, and delivers additional candidate genes to guide future research.

## INTRODUCTION

Food allergy is a significant and increasing public health concern (1,2). As the global burden of food allergy rises, there is a growing need to deepen our understanding of the biological mechanisms driving allergen immune responses. Among food allergies, those against tree nuts are especially concerning due to their severity and association with a reduced quality of life (3,4). Members of the tree nut family include hazelnut, walnut, almond, pecan nut, cashew, pistachio, and Brazil nut. Among these, hazelnut allergy is the most common tree nut allergy in Europe (4), presenting with reactions ranging from mild to life-threatening, particularly amongst children (5–7). Although hazelnut allergy generally manifests in childhood, it typically persists into adulthood, thereby contributing to long-term socioeconomic challenges (8).

Multi-omics studies have successfully identified several mechanisms underlying food allergies, but primarily focused on peanut allergy and peanut oral immunotherapy response (9–11). Few studies have examined the genetic mechanisms underlying tree nut allergy. A genome-wide association study identified a hazelnut-associated locus near *HLA-DPA1*, whereas another study characterised gene expression changes in the absence of antigen-specific immune cell activation (12,13). The role of epigenetics and proteomics in hazelnut allergy also remains unexamined. This highlights the need to further our understanding of hazelnut allergy from a multi-omics antigen-specific perspective.

In this study, we aimed to address these gaps by comparing multi-omics profiles of double-blind placebo-controlled oral food challenge-(OFC) confirmed hazelnut-allergic individuals against nonallergic controls. We integrate in parallel genome-wide methylation profiling, mRNA sequencing, and secreted inflammatory proteins captured from isolated immune cells after 48h stimulation with hazelnut protein. Our findings provide insights into the mechanisms distinguishing hazelnut-allergic from nonallergic individuals, and provide a foundation for further research to inform the development of targeted therapies and improve diagnostic accuracy for hazelnut-allergic individuals.

## METHODS

Detailed methodology is provided in the Supplementary Methods. Software packages and tools are listed in Table E1, and materials used are outlined in Table E2.

### Study cohort

Food allergic and nonallergic individuals were recruited at Charité University Hospital, Berlin, Germany as part of the **T**olerance **I**nduction through **N**on-avoidance to prevent persistent food **A**llergy (TINA) trial (14). Hazelnut-allergic individuals (n = 24) were OFC confirmed and nonallergic individuals (n = 109) defined as having no history of food- or aero-allergies as well as negative skin prick tests. Sample demographics are shown in Table 1.

**Table 1.** Characteristics of the study participants.

| Hazelnut cohort characteristics | Hazelnut Allergic (N = 24)<sup>1</sup> | Nonallergic (N = 109)<sup>1</sup> | P-value<sup>2</sup> |
| --- | --- | --- | --- |
| Age at blood collection (years) | 8 (5, 24) | 26 (22, 30) | <0.001 |
| Sex (F) | 11 (46%) | 70 (64%) | 0.095 |
| History of asthma | 8 (33%) | 2 (1.8%) | <0.001 |
| History of Eczema | 19 (79%) | 2 (1.8%) | <0.001 |
| SPT hazelnut roasted (mm) | 6.00 (3.00, 8.75) | 0.00 (0.00, 0.00) | <0.001 |
| SPT Birch Pollen (mm) | 7.00 (4.00, 8.00) | 0.00 (0.00, 0.00) | <0.001 |
| SPT Timothy Grass (mm) | 7.75 (4.00, 14.00) | 0.00 (0.00, 0.00) | <0.001 |
| Total IgE (kU/L) | 529 (150, 1,314) | 21 (11, 54) | <0.001 |
| sIgE (kU/L) | 20 (8, 72) | NA | NA |
| Cor a 9 (kU/L) | 1 (1, 10) | NA | NA |
| Cor a 14 (kU/L) | 8 (2, 41) | NA | NA |

Table . Demographics characteristics of the hazelnut cohort and nonallergic controls. Key clinical parameters include age, sex, comorbidities, skin prick test (SPT) results, and serum IgE total and specific (sIgE) levels (kU/L = kilo units per litre). NA indicates cases where no data is available.1: In the patient columns, the data is formatted either as the number and percentage of participants in each category or as Median (Q1, Q3). 2: The p-value is calculated by Wilcoxon rank sum test, Pearson's Chi-squared test, or Fischer's exact test.

### Hazelnut extract preparation

Hazelnut protein extract was prepared from roasted hazelnut (Seeberger, Kaufland). Stock protein concentration was 21.3mg/ml (Pierce^TM^ BCA assay). Endotoxin was removed using Pierce™ High-Capacity Endotoxin Removal Resin; diluted aliquots (2.51mg/ml) had endotoxin levels of 0.484EU/ml. At a working concentration (100µg/ml), endotoxin content was 0.02EU/ml, below the accepted FDA limit of 0.5EU/ml for medical devices (15).

### Cell culture

Whole blood was collected into sodium heparin tubes, in allergic individuals before OFC and in nonallergic individuals during screening. Peripheral blood mononuclear cells (PBMCs) were isolated using SepMate^TM^ tubes and BioColl^®^ density gradient centrifugation and cryopreserved. PBMCs were cultured for 48 hours in the absence (quiescent) or presence (stimulated) of 100µg/ml hazelnut protein extract in a total of 500µl media with a seed of 2E+06 cells.

### DNA, RNA, and protein isolation

Following 48-hour cell culture, DNA and RNA were co-extracted from cell pellets using a combination of Qiagen AllPrep DNA/RNA/miRNA Universal kit, Zymo Clean & Concentrator-5 kit, and TRIzol^®^-chloroform protocols. Cell culture supernatant was stored at -80°C until further use. DNA concentration and purity were assessed using a Nanodrop. DNA was stored at -20°C until further use. RNA integrity (RIN) of diluted RNA was quantified using the RNA 6000 pico Bioanalyser kit. All RNA samples presented with a RIN ≥ 8 and were used for mRNA sequencing. RNA was stored at -80°C until further use.

### Profiling and analysis of DNA, RNA and protein data

DNA samples were randomised and processed on the Infinium MethylationEPIC v1.0 array. Raw .idat files were processed in the R environment using Minfi. Two individuals were excluded for sex mismatch and 725,817 probes remained for statistical analysis. Differentially methylated probes (DMPs) were identified using the limma Bioconductor package at a genome-wide significance threshold of 9E-08 (16,17). The Enmix comb-*p* algorithm was used to identify differentially methylated regions (DMRs) (18). Significant DMRs were defined as regions represented by at least three probes and a Šidák-corrected *P* ≤ 0.05. Annotatr was used to annotate DMRs using the hg38 genome as reference (19). Methylation beta-values were converted into M-values for statistical analysis.

Total RNA was processed using the Truseq Stranded mRNA Library Prep Kit. The Spliced Transcripts Alignment to a Reference (STAR) aligner was used to map and count gene transcripts. Differential gene expression was performed using DESeq2 (20). Differentially expressed genes (DEGs) were defined having a false discovery rate (FDR)-adjusted p-value of less than 0.05 (*P_FDR_* ≤ 0.05) as well as an absolute log2 fold change (log2FC) ≥ 0.3.

Thawed cell culture supernatants (40µl) were randomised into 96-well plates and processed using the Olink® Target 96 inflammation panel. After quality control (QC) excluding lowly detected signals, 74 of 92 proteins were retained in the final analysis. Differentially expressed proteins (DEPs) were defined as *P_FDR_* ≤ 0.05.

In total, 218 nonallergic and 46 hazelnut-allergic samples were used in the final analysis. Analyses were performed in R version 4.3.2 (2023-10-31). RNA demultiplexing and mapping were used in the bash environment. Table 2 describes the statistical models used in this data.

**Table 2.**
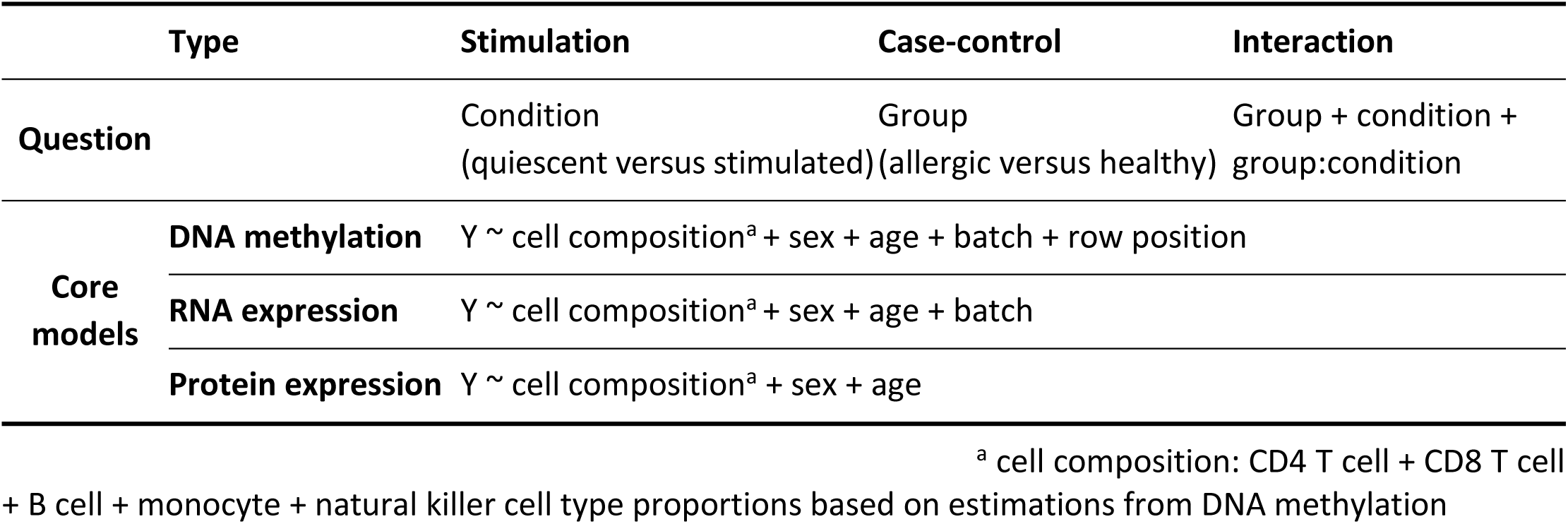
Summary of examined statistical models and comparisons.

<sup>a</sup> cell composition: CD4 T cell + CD8 T cell + B cell + monocyte + natural killer cell type proportions based on estimations from DNA methylation
|  | Type | Stimulation | Case-control | Interaction |
| --- | --- | --- | --- | --- |
| Question |  | Condition<br>(quiescent versus stimulated) | Group<br>(allergic versus healthy) | Group + condition +<br>group:condition |
| Core<br>models | DNA methylation | Y ~ cell composition<sup>a</sup> + sex + age + batch + row position |  |  |
|  | RNA expression | Y ~ cell composition<sup>a</sup> + sex + age + batch |  |  |
|  | Protein expression | Y ~ cell composition<sup>a</sup> + sex + age |  |  |

### eQTM and gene expression protein analysis

Methylation-gene expression Pearson correlation analysis was performed to capture expression quantitative trait methylation signatures (eQTMs). Methylation-DEG correlations with *P_FDR_* ≤ 0.05 were considered significant. EnhancerAtlas 2.0 was used to predict putative promoter and enhancer signatures of DMR coordinates (21). For gene expression-protein Pearson correlations, DEGs and DEPs were overlapped. *P_FDR_* ≤ 0.05 signatures were considered significant. The psych package was used to assess differences in the correlation slopes between allergic and nonallergic groups (*P_r.test_* ≤ 0.05) (22).

### Functional enrichment

Enrichment was performed in R using the gprofiler2 package (23). Significant terms were defined as *P_FDR_* ≤ 0.05, with a minimum query overlap of 4 and term size of 10. Input genes included DMP-/DMR-associated genes (methylation), DEGs (transcriptomics), and DEPs (proteomics). Gene Set Enrichment Analysis (GSEA) was performed using “fgsea” in R, with DEGs ranked by log2FC (24). The human Molecular Signature Database (MSigDB) (v2023.2.Hs) gene sets (term size 15-500) were used; *P_FDR_* ≤ 0.01 was considered significant. Protein-protein interaction (PPI) networks were generated with STRING (25). Clustering was performed with the STRING mcl algorithm using the default inflation parameter of three.

## RESULTS

Study participants were 24 individuals with OFC-confirmed hazelnut allergy and 109 without hazelnut allergy or sensitisation (see Supplementary methods). Baseline characteristics are shown in Table 1. As expected, allergic participants had higher rates of eczema, asthma, pollen and grass sensitisation, and hazelnut-specific IgE, with no significant sex differences between groups.

### Differentially methylated regions with immune relevance detected in hazelnut allergy

DNA methylation was profiled in PBMCs from hazelnut-allergic and nonallergic individuals under quiescent and hazelnut-stimulated conditions. First, we examined individual cytosine-phosphate-guanine (CpG) sites to highlight differentially methylated probes (DMPs). As DNA methylation often spans clusters of neighbouring CpGs, we additionally examined differentially methylated regions (DMRs) to detect broader and more biologically relevant methylation patterns.

Case-control analysis of quiescent samples identified a single hypomethylated DMP (cg07322512; *P* = 7.12E-08; logFC = -0.29; Table E3) in a CpG shore near *SIX3* (Figure 1a). In hazelnut-stimulated samples, a hypomethylated DMP overlapping the first exon of *MFHAS1* (cg03899721, *P* = 6.8E-08; logFC = -0.41) reached genome-wide significance. Additionally, we detected 38 DMRs in quiescent and 46 in stimulated samples (Figure E1/Table E4). Significant DMRs were detected in both analyses, showing modest but consistent methylation differences, with similar effect sides in both conditions (Figure 1b). A significant overlap in DMR-associated genes was observed between quiescent and stimulated samples (29.1%; *P_hyper_* ≤ 0.01), with 22 (27.8%) and 34 (43.0%) genes unique to each condition, respectively (Figure 1c).

**Figure 1.**
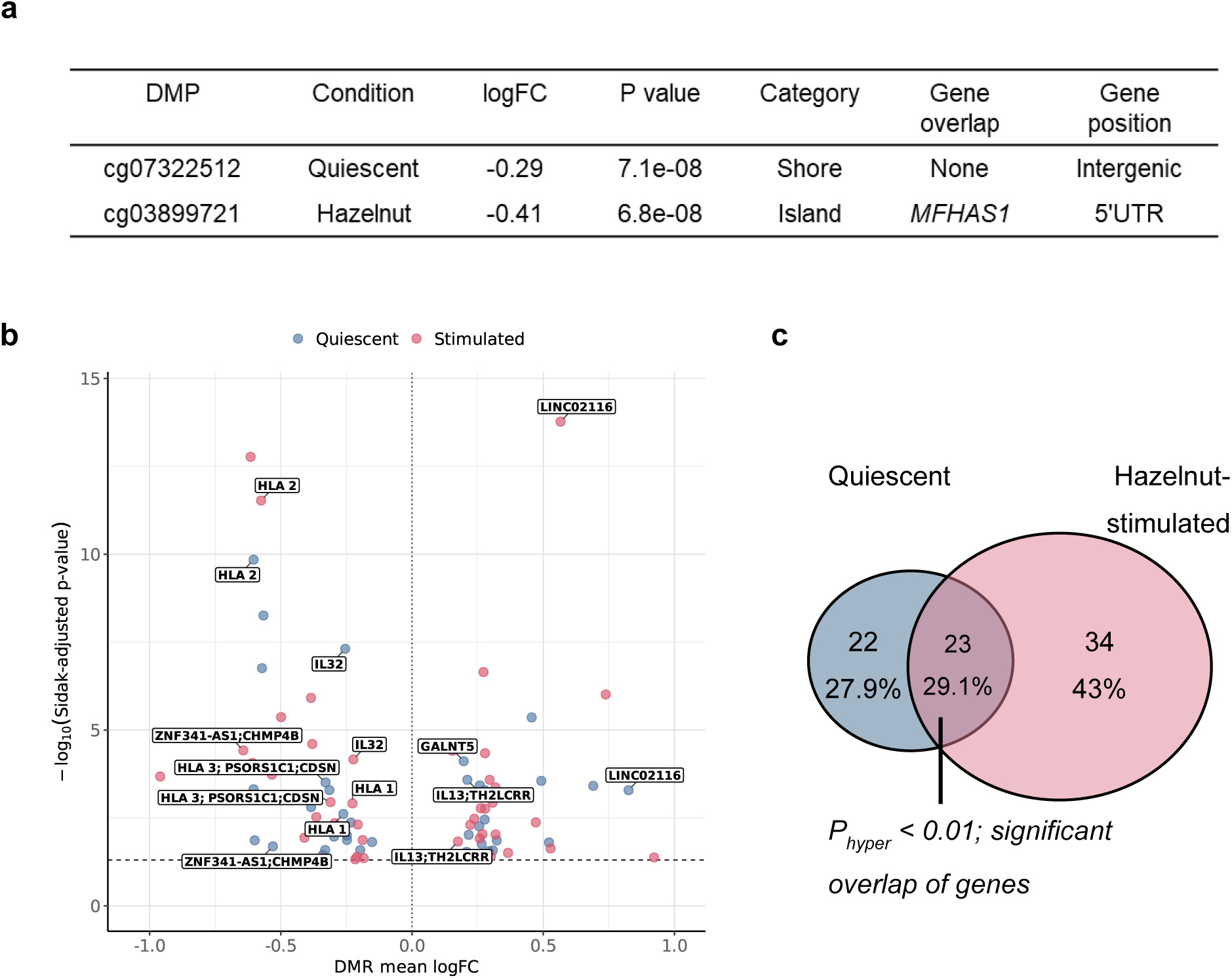
DNA methylation case-control hazelnut overview. (a) Table showing the only two detected DMPs as well as information about these signals. DMP = differentially methylated probe; LogFC = log2 fold change; Category = CpG genomic context. (b) Volcano plot illustrating adjusted p-values and average effect sizes of each DMR, coloured by quiescent (blue) and stimulated (red) analyses. Genes of interested are labelled. (c) Venn diagram depicting the DMR-associated genes and their overlap in quiescent and hazelnut-stimulated PBMCs.

Overlapping genes included *GALNT5* (quiescent), as well as the immune-related *IL13/TH2LCRR*, *IL32*, and *MFHAS1* (shared; Figure 1c, Table E4). In quiescent samples, a hypermethylated DMR overlapped the *GALNT5* promoter and first exon (chr2:157257557-157257913; *P_Šidák_* = 7.72E-05; logFC_µ_ = 0.20).

A DMR spanning the *IL32* promoter to second exon was significantly hypomethylated in hazelnut-allergic versus nonallergic individuals in both quiescent (Figure 1c; chr16:3065285-3065809; logFC = -0.25; *P_Šidák_* = 4.91E-08) and stimulated (chr16:3065551-3065809; logFC = -0.22; *P_Šidák_* = 6.88E-05) samples. We likewise observed hypomethylated DMRs overlapping the first exon of *MFHAS1* (quiescent: chr8:8891026-8891565; logFC = -0.31; *P_Šidák_* = 5.13E-04; stimulated: chr8:8891026-8891834; logFC = -0.33; *P_Šidák_* = 1.32E-07) as well as DMRs overlapping the antisense RNA gene *ZNF341-AS1* (quiescent: chr20:33810217-33810459; logFC = -0.53; *P_Šidák_* = 2.03E-02; hazelnut-stimulated: chr20:33810217-33810459; logFC = -0.64; *P_Šidák_* = 3.82E-05).

Two of the largest and highly significant hypomethylated DMRs in the HLA class I region showed consistent effects in hazelnut-allergic individuals irrespective of stimulation status (chr6:31307774-31308436/chr6:31307490-31308436; logFC ≍-0.58; *P_Šidák_* ≤ 1.43E-10; and chr6:30127359-30127643/chr6:30127210-30127719; logFC ≍ -0.23; *P_Šidák_* ≤ 2.44E-10; Table E4). A third, smaller hypomethylated DMR overlapping *CDSN* was also detected in both conditions (chr6:31120411-31120658; logFC ≍-0.32; *P_Šidák_* ≤ 1.12E-03).

In contrast, a hypermethylated DMR was detected overlapping the first intron of *TH2LCRR* and upstream of *IL13* (chr5:132655799-132656076) in both conditions (quiescent: logFC_µ_ = 0.21, *P_Šidák_* = 2.61E-04; stimulated: logFC_µ_ = 0.17, *P_Šidák_* = 1.48E-02).

To assess group-condition responses to antigen, we performed an interaction analysis and identified one DMP (cg19017420) in a CpG shore upstream of *ARL2* on chromosome 11 (*P* = 7.11E-08, logFC = -0.47). No functional enrichment or transcription factor motif enrichment was observed for DNA methylation-associated genes in any methylation analysis.

### Gene expression analysis after hazelnut stimulation highlights Th2, activation machinery, and vitamin D-related genes

Gene expression of PBMCs revealed 14 differentially expressed genes (DEGs) in quiescent samples and 118 in stimulated samples, most of which were upregulated in allergic individuals (Figure 2a, Table E5). Seven DEGs (5.6%) overlapped analyses (*P_hyper_* ≤ 0.05; significant overlap of genes), while 88.8% were unique to the stimulated state (Figure 2c). Highly up-regulated allergy-related genes included *IGHE*, *PTGDR2*, *IL9*, *IL17RB*, *IL13*, and *IL5* (Figure 2b). Additional stimulation-specific signals included *NSMCE1-DT*, *CSF1/2*, *IL3*, and *CYP27B1*. Top quiescent DEGs included *CYP3A5*, *SYT4*, *ITGB2-AS1*, *SEMA5A, FREM1*, and *HMOX1* (Figure E2).

**Figure 2.**
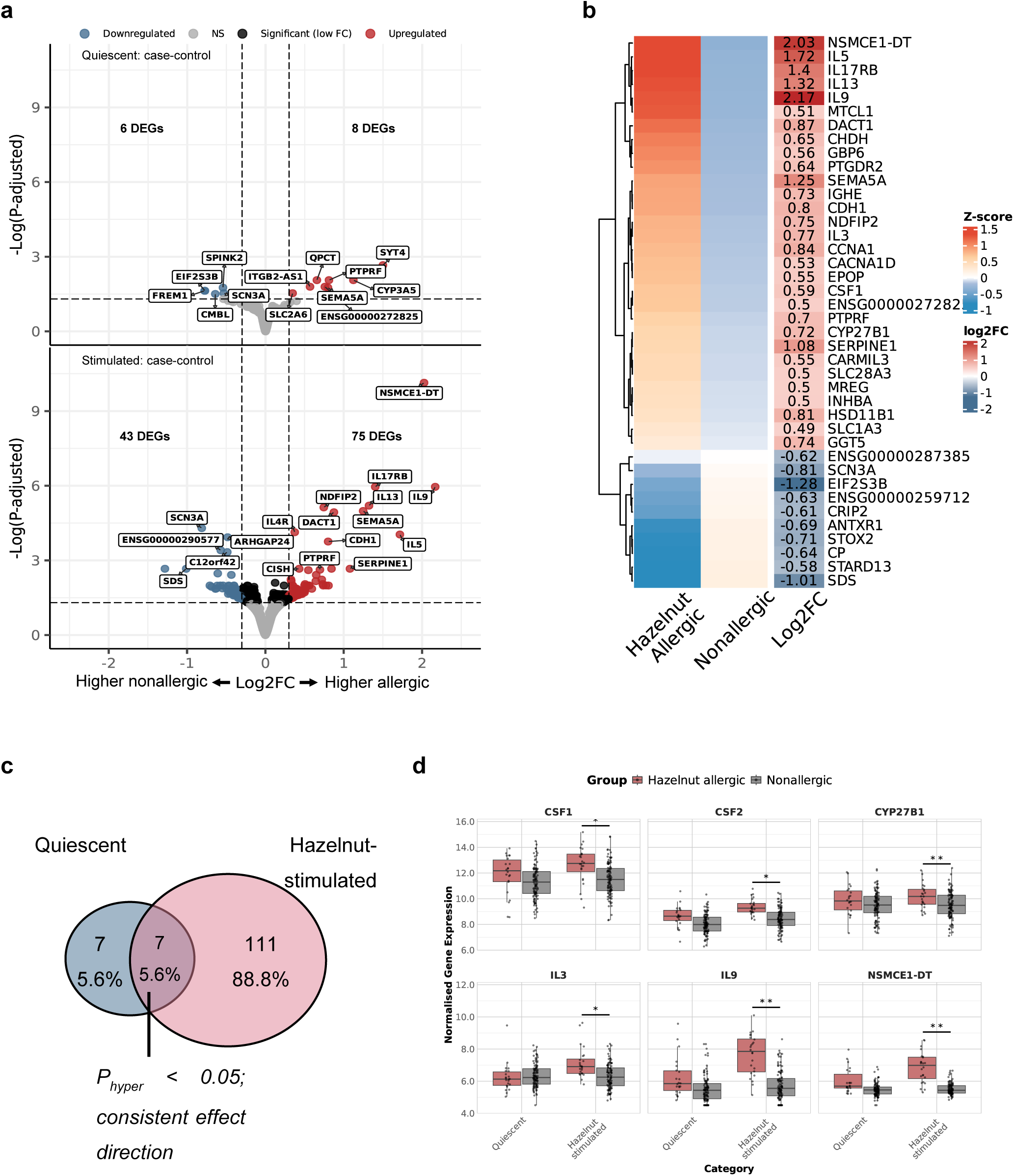
Summary of case-control hazelnut DEGs. (a) Volcano plots depicting the DEGs in quiescent (top) and hazelnut-stimulated (bottom) case-control analysis. An absolute log2FC threshold of 0.3 was applied and highlighted beyond the vertical dotted line. The 20 most highly and 10 least expressed genes are labelled, where possible. (b) Heatmap of average Z-scores for the top DEGs in the stimulated analysis. (c) Venn diagram showing the overlap of DEGs between analyses. (d) Boxplots illustrating a subset of the top immune- or allergy-related DEGs and their significance. Significant genes are marked by asterisks (* = P_FDR_ ≤ 0.05 and ** = P_FDR_ ≤ 0.01). DEG = differentially expressed gene.

To characterise biological processes linked to hazelnut allergy, we assessed enrichment of upregulated and downregulated genes using curated pathways. Key GO biological processes identified in hazelnut stimulated samples included cytokine responses, STAT signalling, immunoglobulin regulation, and cell growth (Figure 3a, Table E6). Up-regulated genes in allergic individuals primarily drove these terms. KEGG analysis pinpointed JAK-STAT, FcεRI, and IL-17 signalling pathways, while Reactome enrichment pointed to interleukin signalling (Figure 3b). Using DEGs associated with hazelnut-allergy, we identified a core protein-protein interaction network of JAK-STAT and interleukin signalling (Figure 3c). JAK-STAT gene contributors included allergy-associated genes (*IL4R*, *IL5*, *IL9*, *IL13*), CSF genes (*IL3*, *CSF2*), as well as *CISH* and *IL2* (Figure E2). GSEA using the Hallmark gene sets highlighted E2F, MYC, G2M checkpoint, oxidative phosphorylation, TNF-α, STAT, and interferon response pathways in up-regulated genes post-stimulation (Figure E2).

**Figure 3.**
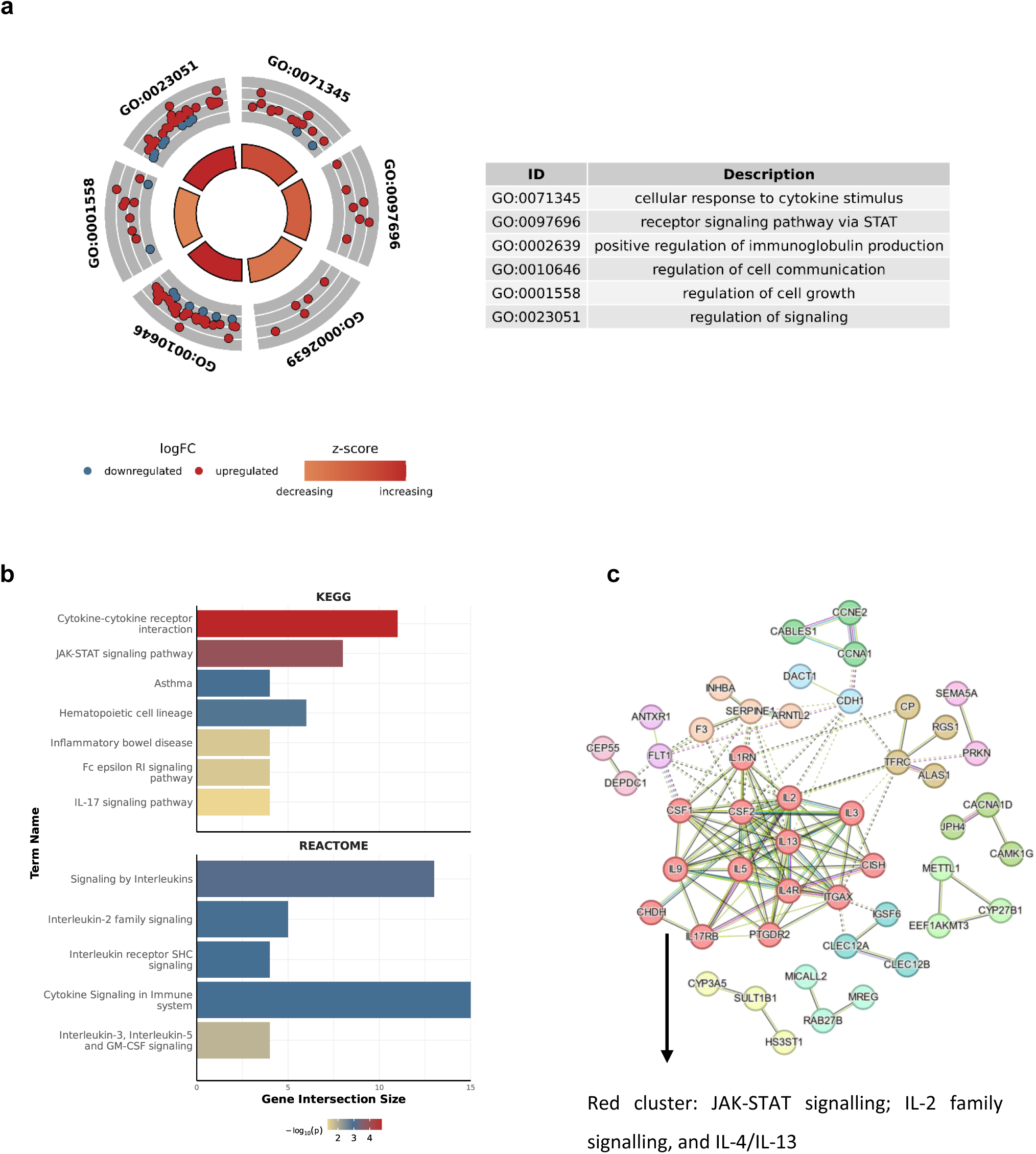
Functional enrichment of genes associated with hazelnut allergy detected after hazelnut stimulation. (a) GOplot summarising the top key driver terms associating with hazelnut allergy. Terms are ordered by their position on the plot axis. Genes contributing to the terms are shown as red (positive logFC/up-regulated) or blue (negative logFC/down-regulated). A z-score indicated the balance of up- versus down-regulated genes contribute to each term. Darker red indicates, for example, that more up-regulated genes contribute to the GO term. (b) All of the KEGG and Reactome terms associating with hazelnut allergy. (c) Protein-protein interaction network of DEGs highlighting a core cluster of genes involved in the JAK-STAT signalling and IL-signalling pathways. logFC = log2 fold change; DEG = differentially expressed gene; GO = Gene Ontology.

### JAK-STAT signalling enrichment in differentially secreted proteins

We analysed differences in protein expression between allergic and nonallergic individuals as a highly functional readout of antigen stimulation. We observed similar numbers of DEPs in quiescent (n = 5) and stimulated (n = 7) samples (Figure 4a; Table E7). VEGFA and IL-12β were detected in both conditions. TNF (TNF-α) was elevated in allergic individuals under quiescent conditions, while TNF-β was increased post-stimulation (Figure 4a/c). KEGG analysis of stimulated samples identified three enriched pathways: “pathways in cancer”, “JAK-STAT signalling” and “inflammatory bowel disease” (Table E8). PPI clustering confirmed modules related to JAK-STAT and inflammatory bowel pathways (Figure 4b), driven by IL-4, IL-13, IL-5, IL-12β, and IFN-γ (Figure 4c). Interaction analysis of the protein data identified five DEPS with significant group-condition effects (*P_FDR_* ≤ 0.05; Figure E3). IL-2, IL-4, IL-5, IL-13, and TNF-β (*LTA*) were all more strongly induced in allergic individuals after antigen stimulation (log2FC = 0.83-1.07; *P_FDR_* ≤ 0.04).

**Figure 4.**
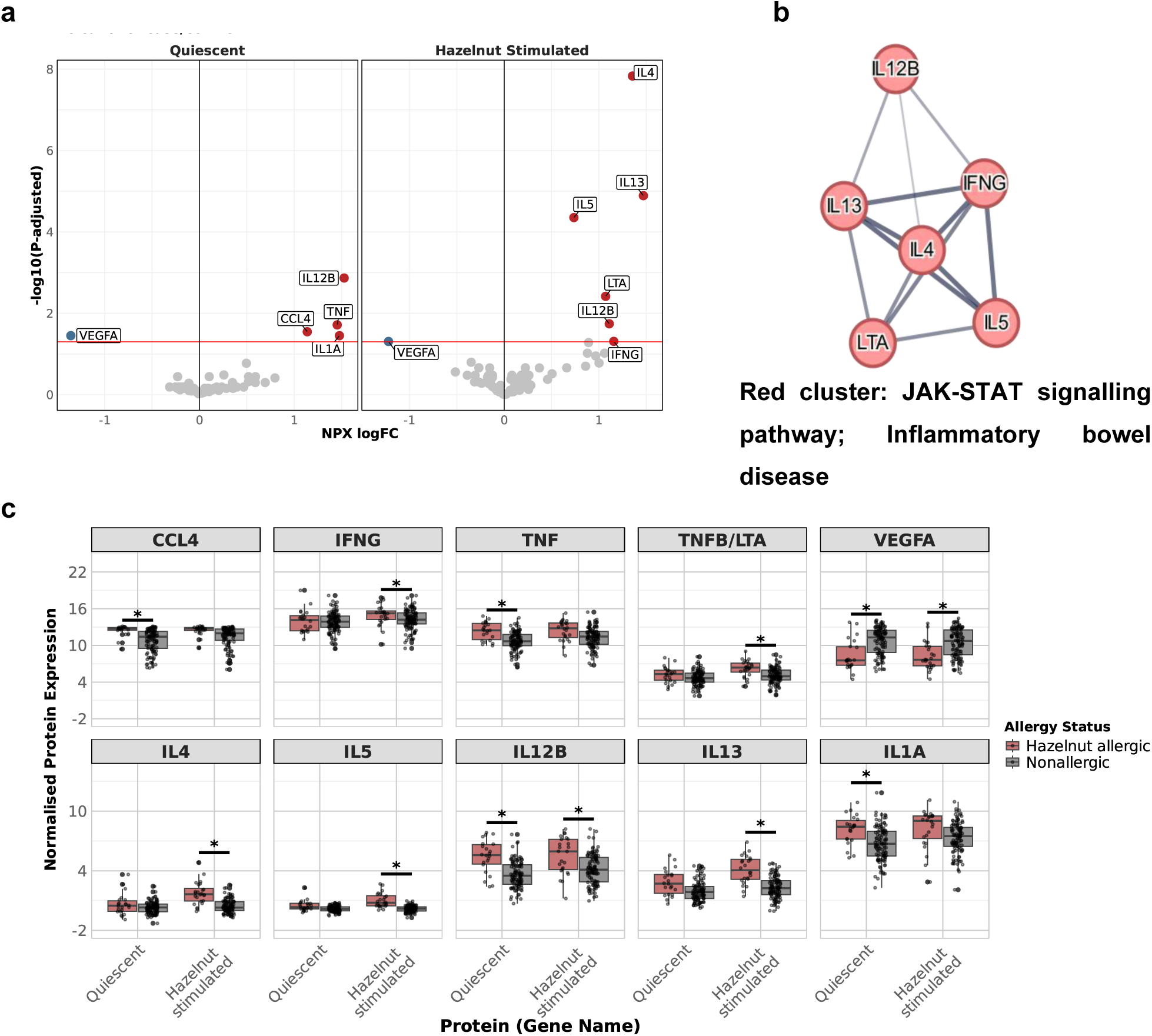
Summary of DEPs detected in hazelnut case-control analysis from 48-hour cell culture supernatant. (a) Volcano plots highlighting all the DEPs found in the case-control quiescent (left) and hazelnut-stimulated (right) analyses. (b) Protein-protein interaction DEPs from hazelnut-stimulated samples showing a cluster involved in JAK-STAT signalling. (c) Boxplots highlighting all DEPs. DEP = differentially expressed protein.

### Multi-omics data integration in hazelnut allergy

We integrated multi-omics data to characterise regulatory differences between hazelnut-allergic and nonallergic individuals under quiescent and antigen-specific conditions. eQTM analysis identified nine DEG-DMR pairs across five DMRs involving *IL5*, *IL13*, *CSF2*, and *IL3*, with *IL5* and *IL13* implicated in each analysis, although none of these correlations remained significant following multiple correction (Figure 5a; Table E9). Similarly, we tested for correlation of gene expression and protein secretion. Two strong DEG-DEP correlations were identified post-stimulation: *IL13* and *IL5* (R > 0.82, *P_FDR_* ≤ 0.001; Figure 5b). Allergic individuals showed a strong correlation for both signals (R > 0.82, *P_FDR_* < 0.001). In nonallergic individuals, *IL13* expression also showed a strong gene-protein correlation (R = 0.77, *P_FDR_* = 2.47E-22), while *IL5* showed a much weaker correlation in these individuals, likely driven by a lack of mRNA detection (R = 0.31, *P_FDR_* = 0.001). Importantly, correlating *IL5* gene expression and protein levels could discriminate between allergic and nonallergic individuals (*P_r.test_* < 0.001), whereas that was not true for *IL13* (*P_r.test_* < 0.57).

**Figure 5.**
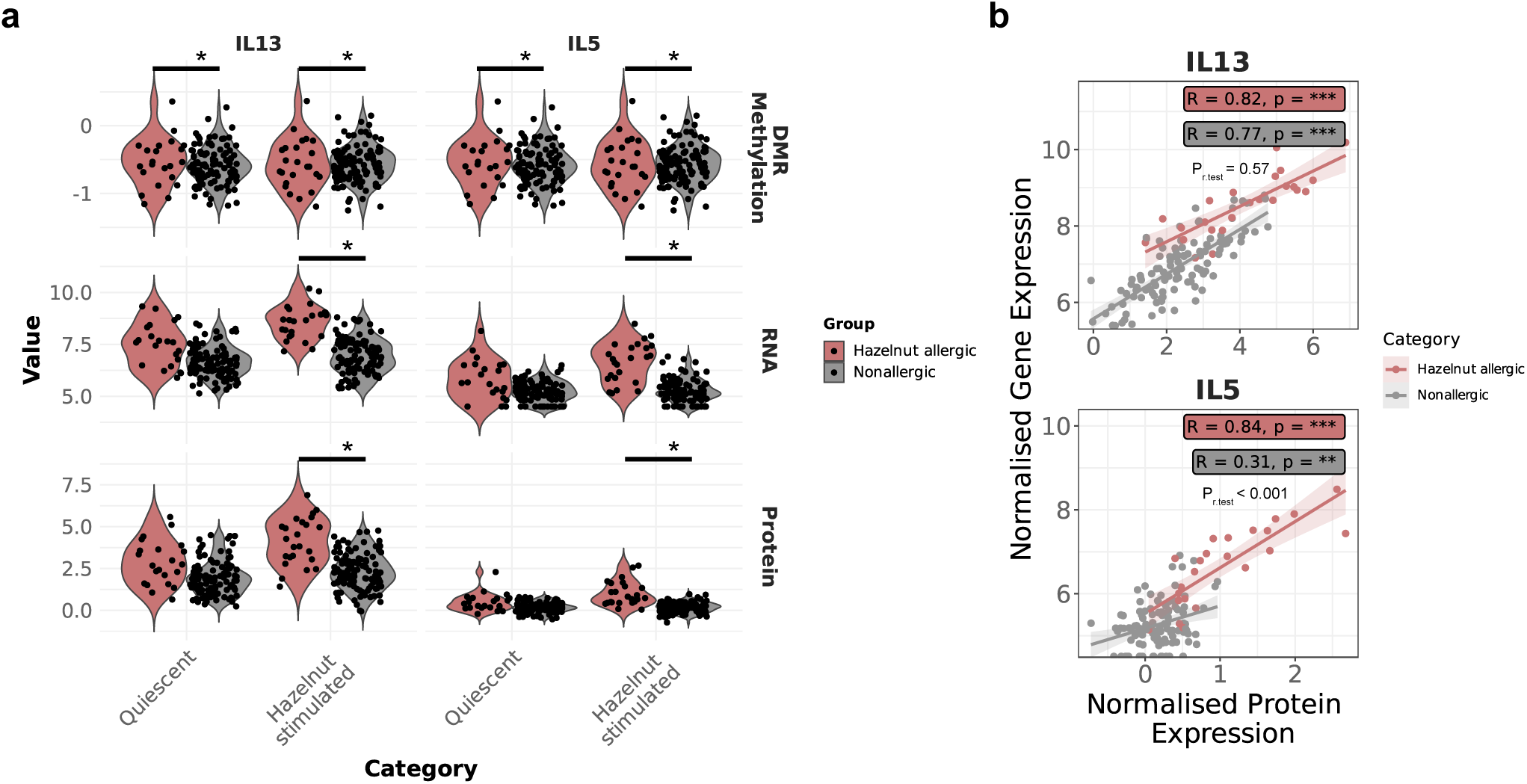
Multi-omics analysis of *IL5* and *IL13* in hazelnut case-control samples. (a) Violin plots showing a single DMR (top) that potentially interacts with *IL5* and/or *IL13* in the gene expression (middle) and protein (bottom) data layers. Significance is denoted by asterisks (P_FDR_ ≤0.05). (b) Pearson correlations of gene expression and protein expression data for *IL5* and *IL13*.

## DISCUSSION

Hazelnut allergy is a major cause for food-related anaphylaxis, yet remains poorly characterised molecularly (26). We therefore used antigen-specific multi-omics profiling to compare immune responses in hazelnut-allergic and nonallergic individuals and identify disease mechanisms and candidate molecular targets.

### STAT3-mediated IgE dysregulation in hazelnut allergy

Epigenetic alterations mediate genetic and environmental effects in complex diseases (27,28), including hazelnut allergy. We found a highly significant overlap between hazelnut allergy-associated DNA methylation patterns both with and without antigen stimulation. This finding confirms the robustness of our technical workflow providing experimental replication of the methylation measurements and is consistent with stable methylation changes underlying long-term effects on gene regulation. In stimulated cells, the top hypomethylated gene-associated DMR overlapped the antisense transcript of *ZNF341*, *ZNF341-AS1*; in quiescent cells, this region remained similarly hypomethylated. Interestingly, homozygous nonsense mutations in *ZNF341*, which encodes a zinc finger transcription factor, have recently be shown to cause autosomal recessive hyper-IgE syndrome. ZNF341 binds to the *STAT3* promoter and is required for its transcriptional activation (29). Notably, *STAT3* mutations cause autosomal dominant hyper-IgE syndrome impairing cytokine signalling, Th17 cell development, and epithelial repair (30), and often featuring food allergy (31). We detected another hypomethylation signal implicating *STAT3* dysfunction, a DMP overlapping the *ARL2* promoter, a gene with diverse functions that is also required for nuclear retention of activated *STAT3* (32). Our results link epigenetic alterations at the *ZNF341* and *ARL2* locus with STAT3-mediated IgE dysregulation in hazelnut allergy (33).

### Methylation of Th2 cytokine control locus

We identified a hypermethylated DMR overlapping the first exon of the *long non-coding T Helper Type 2 Locus Control Region Associated RNA* (*TH2LCRR*) which is known to upregulate the expression of Th2 cytokines *IL4*, *IL5*, and *IL13* (34,35). The DMR is located 14 kb upstream of the dominant *TH2LCRR* isoform in immune cells (transcript ENST00000457489.1, exons 4-7, GTEx (36)). While hypermethylation is mostly associated with decreased gene expression, a recent genomewide study in whole blood found that it is associated with increased expression in 34% of cases (37). Our findings suggest that hypermethylation of this DMR may mediate the upregulation of IL-4, IL-5 and IL-13 cytokine expression which we also detected.

Furthermore, integration of RNA and protein data revealed that *IL5*, but not *IL13*, was able to discriminate between allergic and nonallergic individuals. Further studies are required to evaluate its potential as disease-specific biomarker. Our data emphasise the importance of antigen-specific stimulation for its detection.

Enrichment of differentially expressed genes highlighted canonical pathways in hazelnut allergy: FcεRI signalling, type-2 cytokine signalling (IL-4, IL-5, IL-13), and the JAK–STAT pathway. These pathways are linked with FcεRI activation releasing Th2 cytokines which signal through the JAK–STAT pathway, thus bridging the immediate effector response to the longer-term transcriptional programs that sustain and amplify disease. This underscores the IL-4/IL-13–JAK/STAT axis as a therapeutic target. Notably, JAK1/2 inhibitors are approved for atopic dermatitis and asthma (38). and a phase I pilot trial is testing the JAK1 inhibitor abrocitinib in food allergy (NCT05069831).

### Three HLA-associated methylation signals

The HLA region, central to antigen presentation, is frequently associated with food allergy. In hazelnut-allergic individuals, we identified three hypomethylated DMRs in the HLA class I region, all present with and without hazelnut stimulation and displaying similar effect sizes within conditions. The first lies between *TRIM31* and *TRIM40*, ubiquitin ligases involved in antiviral immunity and excessive inflammation response (39). Interestingly, upregulated intestinal *TRIM40* expression has been found in chronic inflammatory bowel disease and intestinal epithelial barrier deficiency which may also play a role in hazelnut allergy (40). The second overlaps *CDSN* (corneodesmosin), where homozygous loss has been associated with severe skin barrier deficiency and food allergy (41). Thus, our study identifies epithelial barrier genes of the gut and skin, extending previous findings on the genetic risk for hazelnut allergy conferred by loss-of-function mutations in the skin barrier gene filaggrin (42). The third lies between HLA-B and HLA-C, genes not previously linked to food allergy, and is the closest DMR (∼1.7 Mbp) to a known hazelnut-associated genetic locus (13).

### Vitamin D Pathway Activation in Hazelnut Allergy

*CYP27B1* (*25-hydroxyvitamin D3 1-alpha-hydroxylase*), which converts calcidiol [25(OH)D3] to the biologically active form calcitriol [1,25(OH)2D3], was upregulated upon stimulation in allergic individuals (43). Interestingly, the role of vitamin D is further supported by the hypomethylation of a DMR spanning promotor and first exon of *IL32*. IL-32 up-regulates CYP27B1 and vitamin D receptor (VDR) expression (44). The complex role of vitamin D in allergic disease has been widely investigated, with both deficiency (45) and excess (46–48) of vitamin D exposure associated with an increased risk of allergy. Interestingly, a randomised, controlled trial of daily vitamin D supplementation in the first year of life, found that higher vitamin D levels in cord blood were associated with higher risk of food allergen sensitisation. Moreover, high-dose vitamin D supplementation was associated with an increased risk of milk allergy (49). Our findings also indicate an upregulation of the vitamin D pathway in hazelnut allergy, although targeted studies examining the pro-inflammatory nature of this response are warranted. To our knowledge, this is the first molecular evidence implicating vitamin D in hazelnut allergy.

### Emerging Epigenetic and Transcriptomic Signals in Hazelnut Allergy

Additional methylation and expression signals point to new hazelnut allergy biology. DMRs at *PHACTR1*, *MFHAS1*, and *SPRED2* replicate findings from peanut-allergy studies (50,51), even though their role in food allergy remains unknown. In resting cells, the promoter/first exon of *GALNT5*, an O-glycosylation enzyme, was hypomethylated. Along with previous EWAS associations of *GALNTL4* (51,52), our findings extend interest in the GalNAc-T gene family in food allergy.

Transcriptomically, the most induced transcript after stimulation in hazelnut-allergic individuals was the lncRNA *NSMCE1-DT*; variants at *NSMCE1* associate with eczema and asthma (53,54). Although the functions of these signals remain undefined, they highlight plausible contributors to hazelnut allergic responses and identify candidates for further investigation.

### Strengths, limitations, and conclusions

No previous work has examined hazelnut allergy using DNA methylation or multi-omics approaches. Strengths of this study include a well-phenotyped study cohort, a strict case definition of OFC-confirmed primary hazelnut allergy, the use of antigen-specific stimulation, strict QC pipelines, and rigorous correction for multiple testing. The absence of genomic inflation, concordant signals across conditions, and recovery of known allergy genes and proteins support the data robustness. Moreover, this data demonstrated that antigen stimulation was required to reveal meaningful hazelnut-specific transcriptional and proteomic signals, but the same was not true for DNA methylation, aligning with long-term stable epigenetic remodelling.

Limitations include the use of PBMCs (limiting cell-specific resolution), the age imbalance between groups, and a single 48-hour timepoint which will have missed dynamic regulatory changes. A small sample size likely resulted in a limited detection of subtle group-specific effects and interactions. Since age-matched controls were not available, we mitigated age effects using statistical modelling. While our sample size did not allow for an independent replication, future studies with larger cohorts will be essential to confirm and extend these findings.

In summary, we generated a detailed register of differentially expressed and/or methylated genes associated with hazelnut allergy. Beyond replicating well established genes and pathways such as the type-2 cytokines, FcεRI signalling and the JAK–STAT pathway, our study pinpoints several novel disease associations that link hazelnut allergy with monogenic hyper-IgE syndromes, emphasise an activation of the vitamin D pathway, and identify novel intestinal and cutaneous epithelial barrier gene in hazelnut allergy. Despite limitations, this study represents the largest and only known hazelnut allergic and nonallergic cohort to examine antigen-specific responses from a multi-omics perspective. Altogether, we provide a catalogue of candidate biomarkers for hazelnut allergy to guide mechanistic studies and prospective validation.

## Acknowledgements

We would like to thank Matthias Ziehm from the Proteomics facility at the Max Delbrück Center, Berlin-Buch, for measuring Olink® samples. We also thank Nadine Fricker for assistance with DNA profiling, and the Genomics facility at the Max Delbrück Center Berlin for sequencing the RNA samples. We are grateful to Susanne Blachut and Claudia Langnick for their support on RNA library preparation. We thank the Deutsche Forschungsgemeinschaft (DFG, German Research Foundation) for funding the clinical research unit (CRU339); Food allergy and tolerance (FOOD@)—409525714.

## LIST OF ABBREVIATIONS

**Abbreviation Description**
BMIQ: Beta-MIxture Quantile dilation
CSF: Colony-stimulating factors
DEG: Differentially expressed gene
DEP: Differentially expressed protein
DMP: Differentially methylated probe
DMR: Differentially methylated region
DMSO: Dimethyl sulfoxide
eQTM: Expression quantitative trait methylation signature
EWAS: Epigenome-wide association study
FBS: Foetal bovine serum
GSEA: Gene Set Enrichment Analysis
IDOL: Identifying Optimal Libraries
logFC/log2FC: log2 fold change
MSigDB: Molecular Signature Database
OFC: Oral food challenge
PBMC: Peripheral blood mononuclear cell
PCA: Principle component analysis
PPI: Protein-protein interaction
QC: Quality control
RIN RNA: integrity
RPM: Revolutions per minute
SNP: Single nucleotide polymorphism
SPT: Skin prick test
ssNOOB: Single-sample normal-exponential out-of-band
STAR: Spliced Transcripts Alignment to a Reference
TINA: Tolerance induction through non-avoidance to prevent persistent food allergy

