## Supplementary Figures for "Multi-omics analysis reveals vitamin D metabolism, hyper-IgE genes, and epithelial barrier dysfunction in hazelnut allergy"

### Slide 1
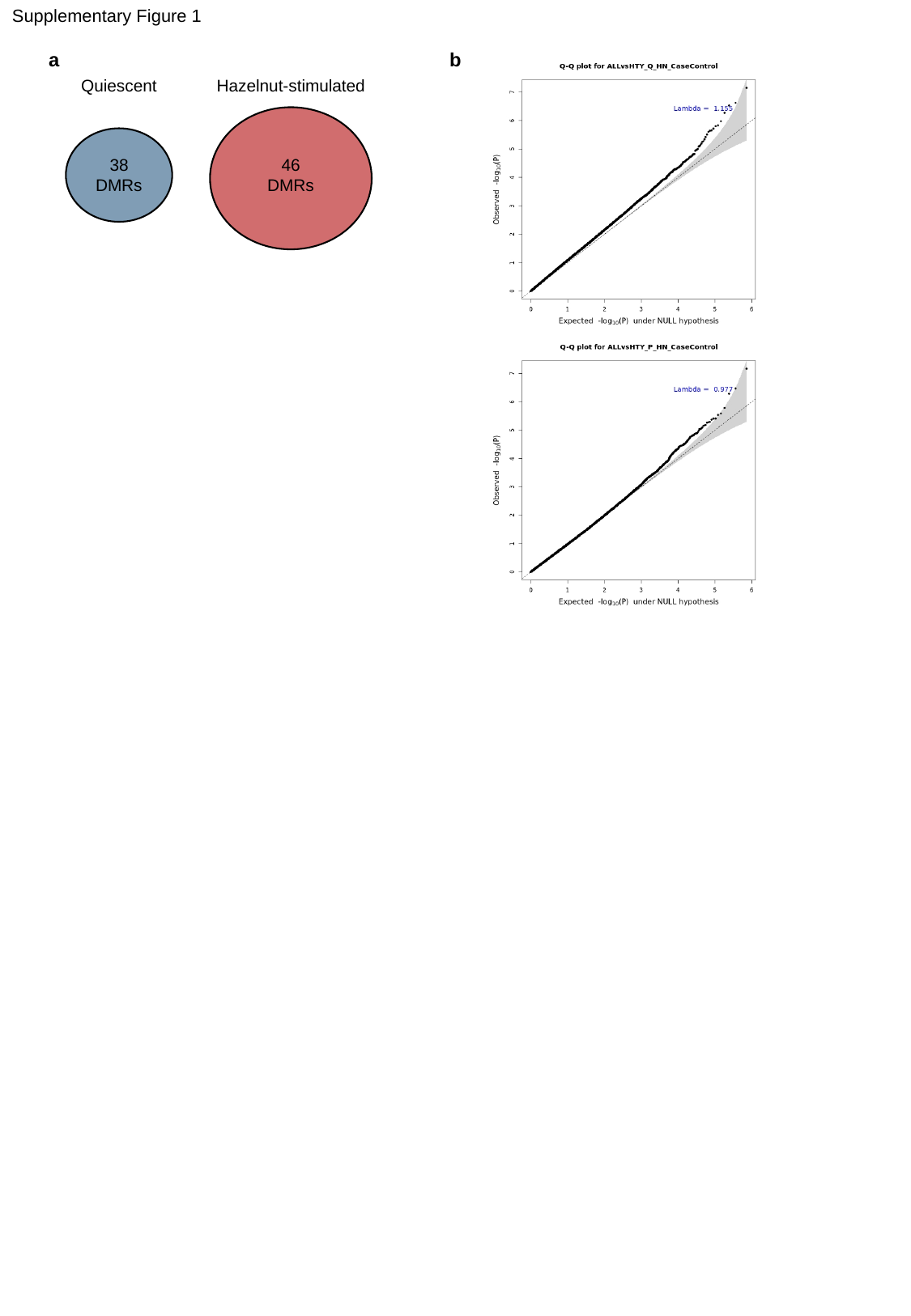

Supplementary Figure 1
a
b
Quiescent
38
DMRs
Hazelnut-stimulated
46
DMRs

### Slide 2
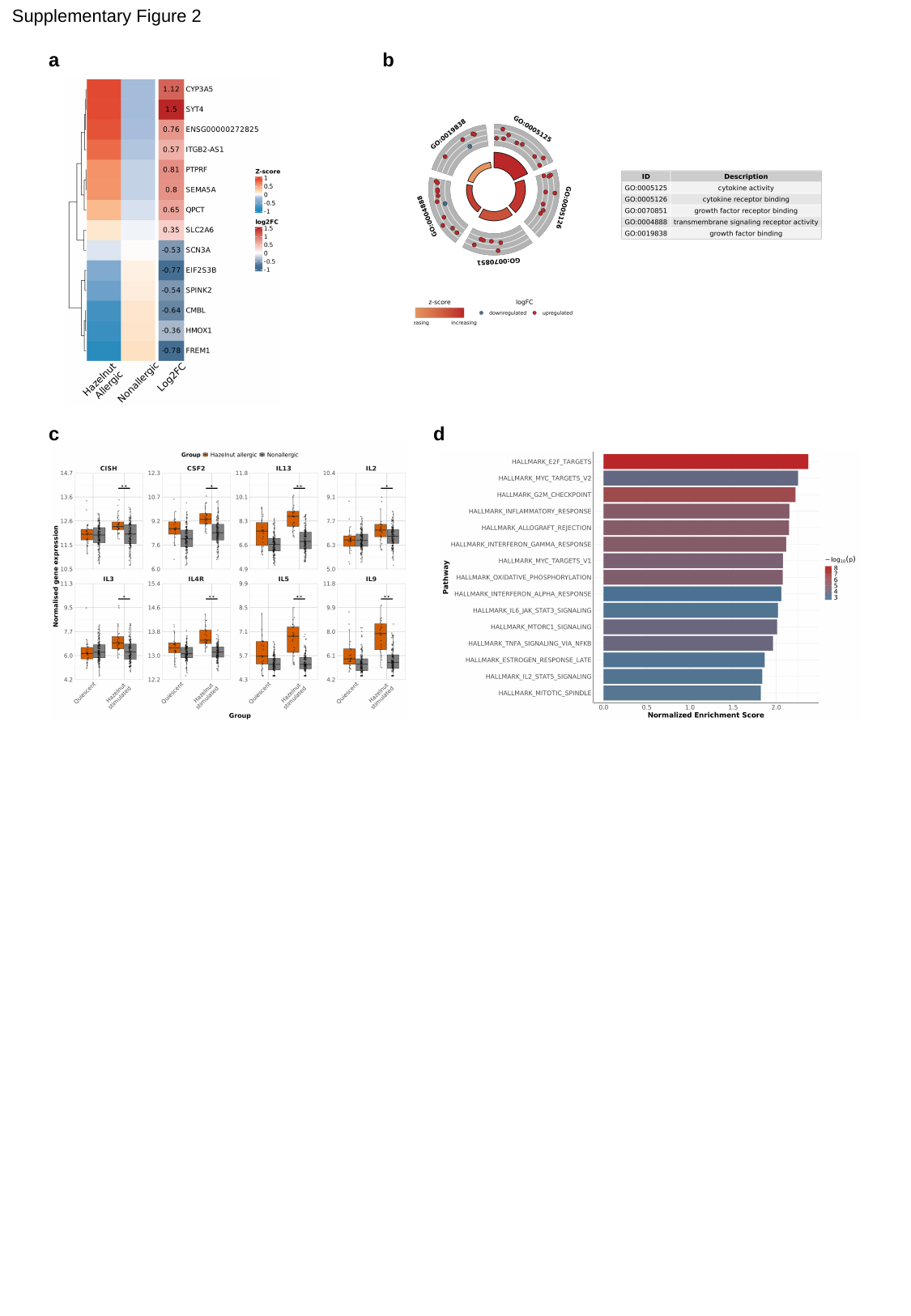

Supplementary Figure 2
a
b
c
d

### Slide 3
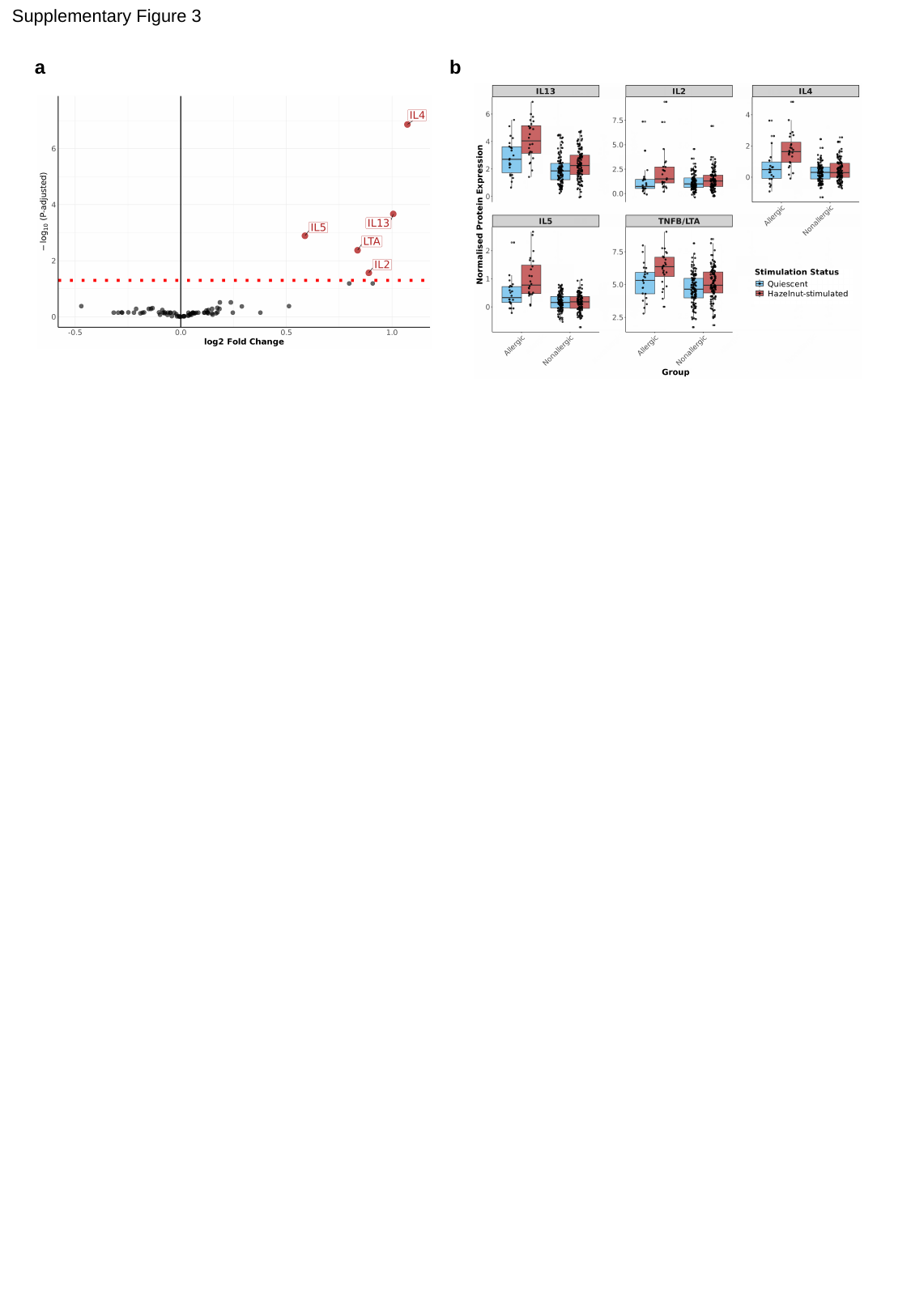

Supplementary Figure 3
a
b
