## Supplementary Methods for "Multi-omics analysis reveals vitamin D metabolism, hyper-IgE genes, and epithelial barrier dysfunction in hazelnut allergy"

**Supplementary Materials**

### Study cohort, extract preparation and cell culture

Samples from individuals in this study were approved for collection under the following ID: trial Ethics No. EA2/033/19 and EA2/304/21; German Clinical Trials Register ID: DRKS00016764 (RCT) DRK S00020467 (1). OFC threshold for hazelnut-allergic individuals were defined as reacting to cumulative dose of less than 3058mg hazelnut protein. Nonallergic individuals have had a SPT < 3mm wheal size to peanut (raw/roasted), hazelnut (raw/roasted), cashew (raw), walnut (raw), wheat flour, gluten, birch, mugwort, timothy grass, house dust mite, and cat epithelium.

Hazelnut protein extract was prepared by shredding into flour, followed by 2-hour acetone degreasing (Merck Millipore, SupraSolv), filtration (Macherey Nagel, MN 615), and overnight air-drying. The dried powder was re-suspended in PBS (Corning) with protease inhibitors (Roche), mixed overnight at 4°C, and centrifuged (20min, 2300xg). The lipid layer was discarded, and the supernatant was aliquoted and stored at -20°C.

Following density gradient centrifugation, PBMCs were cryopreserved in liquid nitrogen using a solution of RPMI 1640 supplemented with 10% foetal bovine serum (FBS) and 10% Dimethyl sulfoxide (DMSO). For culture, PBMCs were thawed into RPMI with 10% AB serum and 10% FBS, adjusted to a concentration of 10E+06 cells/ml, with 2E+06 cells suspended in 200µl media cultured in 5ml round-bottom tubes. Quiescent samples received additional 300µl media while stimulated samples received hazelnut protein extract at 100µg/ml in 300µl media. The final cell culture volume was 500µl. If cell yield was limiting, stimulated conditions were prioritised. Cultures were incubated at 37°C with 5% $CO_2$ for 48 hours.

### DNA and RNA isolation

All steps were performed at room temperature, unless otherwise specified. Cell pellets were lysed with syringes in an RLT/1% β-mercaptoethanol buffer, and lysates passed through Qiagen DNA columns. All centrifugation steps occur at 20,000xg for 30 seconds unless otherwise specified. Lysate was centrifuged and the flow-through used for parallel RNA isolation. Bound DNA was washed using 350µl AW1 buffer. Protein was denatured by incubation with 80µl Proteinase K master mix followed by washing with 350µl AW1. 500µl AW2 was incubated with DNA for 4 minutes followed by a 2-minute centrifugation and aspiration of any remaining liquid from the columns. DNA was eluted three times using elution buffer. For each elution, buffer was pre-incubated with DNA for 1 minute followed by centrifugation at 10,000xg for 1 minute. The first two elutions used 50µl buffer while the third used 100µl buffer. For the third elution, buffer-DNA solution was pipetted back into the DNA column and eluted a fourth time. After each elution, samples were immediately transferred to ice.

Flow-through from the first step of DNA isolation was used for RNA extraction. The volume of flow-through was measured followed by the addition of 1:1 chloroform and 3:1 TRIzol®. This solution was mixed vigorously by hand for 1 minute, followed by end-over-end turning on a Hulu Mixer for 9 minutes (RPM 27/10; deg 51°/10; 5°/2). Samples were centrifuged at 12,000xg/4°C for 15 minutes followed by aspiration of the aqueous phase. 1:1 volume of pure molecular-grade ethanol was added to this aspirated solution. RNA was bound to an RNA column from the Zymo Clean & Concentrator-5 kit by centrifugation at 12,000xg for 30 seconds. Bound RNA was washed as per the manufacturers protocol. The single 50µl elutions were placed directly onto ice. Diluted aliquots were made to bring RNA concentration into range for the pico Bioanalyser assay.

**DNA, RNA, and protein profiling and analysis**

DNA was processed across two batches over a two-year period on the Infinium MethylationEPIC v1.0 array, following Illumina's "Infinium HD Methylation Assay Guide" at the Department of Genomics, Life and Brain GmbH in Bonn, Germany. To mitigate batch effects and technical effects for each batch, DNA samples were randomised so that, where possible, each BeadChip contained all paired samples from a single individual (i.e., quiescent and stimulated). Additionally, each BeadChip needed to contain at least one sample from each phenotype (allergic or nonallergic) and broad age group (adult ≥ 18 or child <18). Sample row order was additionally randomised. Lastly, care was taken to ensure as many different patient's phenotypes as possible were represented across each batch of data. Quality control was carried out in parallel with peanut samples accrued from the same study to further identify and mitigate technical batch effects. Probes failing a detection p-value ($P \leq 0.01$) in more than 5% of samples were excluded (n = 1,923). Additionally, probes represented by less than three beads in more than 5% samples were excluded (n = 13,433). Study-specific "bad" probes consisted of 14,999 unique probes. In addition, a list of known problematic probes (cross-reactive, blacklisted, general masked probes, probes with single nucleotide polymorphisms (SNPs) < 5bp from interrogation sites, and XY probes) were excluded (n = 124,686; (2–4)). Overall, 140,042 unique probes were excluded, leaving 725,817 probes in the final analysis. Two nonallergic patients were excluded on the bases of sex mismatch. No samples were excluded on the basis of more than 5% probes failing a detection p-value, having a low bead count, outliers from median methylation vs unmethylation log2 intensity, OOB outliers, clustering using principal component analysis (PCA), or any other quality control metric. 218 nonallergic and 46 hazelnut-allergic samples were used in the final analysis. Data was normalised using a combination of single-sample normal-exponential out-of-band (ssNOOB; dyeCorr = TRUE, dyeMethod = “single”) and Beta-MIxture Quantile dilation (BMIQ; (5–10)). Cell compositions were estimated using the Identifying Optimal Libraries (IDOL) algorithm with the estimateCellCounts2 function for CD4 T cells, CD8 T cells, NK, B cells, and monocytes (11,12). The parameters used to identify DMRs were the following: combp(dmr_list[[i]], dist.cutoff=1000, bin.size=310, seed=0.05, region_plot=TRUE, mht_plot=TRUE, nCores=1, verbose=TRUE). The DNA manifest file used in this study was created using various sources. The primary hg38 manifest was downloaded from http://zwdzwd.github.io/InfiniumAnnotation (also see: https://github.com/zhou-lab/InfiniumAnnotation and https://rpubs.com/zhouwanding/417953). This manifest was created by Zhou et al. 2017 (2). In addition to this, data from various sources were merged with this table (13–15). Genomic inflation remained low in all analyses ($\lambda$ = 1.16 and 0.98; Figure E1).

Total RNA was processed using the Truseq Stranded mRNA Library Prep Kit with 96 RNA UD Indexes v2. Samples were randomised in an identical manner to the DNA methylation and prepared in 96-well plates in parallel with peanut samples of the same study. Libraries were sequenced on the NovaSeq 6000 S4 (batch 1) and NovaSeq X Plus (batch 2). Samples were sequenced at a depth of 50mln 100bp paired-end reads per library. Raw data was converted to FastQ format using bcl2fastq and the reads aligned to the ensemble v109 hg38 genome using the STAR aligner by decompressing paired-end FASTQ reads and aligning in two-pass mode (--twopassMode Basic) and the output counts used as a raw count matrix for further downstream processing (16). FastQC, MultiQC, and PCA were used to evaluate the sample quality. Nonconverging genes in the DESeq2 analysis were excluded from further analysis. Contrasts were extracted using the “ashr” adaptive shrinkage estimator (17).

Protein profiling was carried out using Olink® Target 96 inflammation panel. Randomising took into account balancing for age, sex, and clinical group, while ensuing where possible that paired samples remained within the same plate. Supernatant data were processed in one batch and in parallel with peanut samples from the same study. Overall, 75% (69/92) proteins were detected in over 75% samples. Proteins detected in less than 75% samples were evaluated on a case-by-case basis for inclusion based on group-specific expression patterns. All supernatant processing and protein measurements were carried out by the Proteomics facility headed by Dr Philipp Mertins at the Max Delbrück Center, Berlin-Buch. Protein values were Intensity Normalized (v.2) by the proteomic facility.

**eQTM and gene expression-protein analysis**

For the eQTM analysis, only DMPs or CpGs of DMRs were used as input from the methylation data. From the gene expression data, only DEGs were considered in the final analysis. As methylation signatures can act over large distances, gene start and end boundaries were extended by ± 1Mb. Methylation coordinates were overlapped with extended gene boundaries to capture distant overlapping signals. Genes were filtered by only those defined as DEGs. A Pearson correlation was then performed. P-values were FDR corrected and signatures using a $P_{FDR}$ ≤ 0.05 significance threshold were reported. In order to concatenate DMR signatures, the mean M-values of all probes within each DMR were calculated. M-Values were converted to β-values to average methylation data before being converted back to M-values for statistical analysis. Protein IDs were converted to gene IDs using data provided from Olink's® Target 96 inflammatory panel for visualisation purposes (https://olink.com/products/olink-target-96). For the gene-protein correlations, overlapping DEG and DEP IDs were considered for Pearson correlation. Correlations with $P_{FDR}$ ≤ 0.05 were reported as significant.

**Functional enrichment**

For the input of genes into functional enrichment tools, duplicate gene entries (as is the case with methylation) were collapsed, selecting the most significant hits where applicable. The "custom_annotated" domain scope was used to reduce bias from each of the omics analyses. Background gene sets included: post-QC probes (methylation), post-QC gene symbols (gene expression), and 74 QC-passed proteins (Olink®). PPI networks used the same background gene list strategies. Full PPI networks were built using the default confidence (0.4) and 5% FDR stringency, including experimental, database, co-expression, and text-mining sources. Neighbourhood, gene fusion, and co-occurrence evidence were excluded. Disconnected nodes were hidden.

**Statistics, modelling, and additional analyses**

In Limma, for multi-level paired analyses, individual ID was included as a random effect in the form of the "duplicatecorrelation()" function. To discriminate between Limma and DESeq2 analyses, "LogFC" was used for limma outputs while "Log2FC" was used for DESeq2 outputs, as extracted from their default summary statistics columns. These can, however, be interpreted similarly.

No genome-wide methylation studies have been published for hazelnut allergy. A reference list of gene-associated methylation signals was compiled from existing peanut-focussed EWASs (18–20). Overlap with current data was assessed.

Plots were generated primarily in the R statistical environment using the "ggplot2", "ComplexHeatmap", "GOplot", "rrvgo", "pathview", or "Gviz" packages (21–26). STRING network plots were exported from the web-interface. Demographics data were tabulated using "gtsummary" in R (27). "PCAtools" was used to calculate principal components and correlate these to clinical variables during QC (28).
